# Size matters: An analytical study on the role of tissue size in spatiotemporal distribution of morphogens unveils a transition between different Reaction-Diffusion regimes

**DOI:** 10.1101/2021.02.16.431401

**Authors:** Alberto S. Ceccarelli, Augusto Borges, Osvaldo Chara

## Abstract

The reaction-diffusion model constitutes one of the most influential mathematical models to study distribution of morphogens in tissues. Despite its widespread use, the effect of finite tissue size on model-predicted spatiotemporal morphogen distributions has not been completely elucidated. In this study, we analytically investigated the spatiotemporal distributions of morphogens predicted by a reaction-diffusion model in a finite 1D domain, as a proxy for a biological tissue, and compared it with the solution of the infinite-domain model. We explored the reduced parameter, the tissue length in units of a characteristic reaction-diffusion length, and identified two reaction-diffusion regimes separated by a crossover tissue size estimated in ∼3.3 characteristic reaction-diffusion lengths. While above this crossover the infinite-domain model constitutes a good approximation, it breaks below this crossover, whereas the finite-domain model faithfully describes the entire parameter space. We evaluated whether the infinite-domain model renders accurate estimations of diffusion coefficients when fitted to finite spatial profiles, a procedure typically followed in Fluorescence Recovery After Photobleaching (FRAP) experiments. We found that the infinite-domain model overestimates diffusion coefficients when the domain is smaller than the crossover tissue size. Thus, the crossover tissue size may be instrumental in selecting the suitable reaction-diffusion model to study tissue morphogenesis.

## Introduction

In their transition towards maturity, tissues are crucially regulated by molecules known as morphogens, whose precise spatiotemporal distribution triggers the downstream changes in protein expression responsible for the exact differentiation patterns. Nevertheless, tissues are not only an inert scaffold upon which morphogens spread, but they are also fully responsible for the morphogen uptake or their transformation by means of specific biochemical reactions. The problem of how a morphogen propagates over a tissue while it is being eliminated was mathematically encoded in the exquisite reaction-diffusion model by the great Alan Turing, who coined the “morphogen” neologism to illustrate its character of “form generator” [1].

The reaction-diffusion model constitutes one of the most influential quantitative approaches within developmental biology. From the aforementioned Turing’s seminal article and the study from Gierer and Meinhardt [2], a progressive wealth of reaction-diffusion models were developed, paving the way to become an essential and pivotal concept to understand tissue morphogenesis [3,4,5,6]. The model was extensively used to investigate distributions of morphogens in a variety of tissues and organisms such as *Drosophila melanogaster* wing imaginal disc [7], chick limb [8] and the stripe pattern of *Danio rerio* [9] among other examples.

Previous studies have analytically investigated this model assuming an infinite domain [10,11]. Although the model relied on the idea that the reaction-diffusion characteristic length of the morphogen pattern was reasonably smaller than the domain, it is clear that biological tissues do not entail infinite lengths. Other reports investigated the model assuming a finite domain by using numerical [7,12] and analytical approaches [13,14,15,16]. To our knowledge, the role played by the size of the domain in the spatiotemporal patterning predicted by this model has not yet been elucidated.

In this study, we present the analytical solution of a reaction-diffusion model describing *de novo* formation of a morphogen and its spread within a finite domain, as a proxy for a tissue. We analytically investigated the behaviour of the model, in terms of a reduced parameter, representing the tissue length in units of a characteristic reaction-diffusion length. We fully characterized the finite-domain model in terms of morphological aspects of the spatial distributions and the time to reach the steady state to finally compare them with the corresponding predictions from the infinite-domain model. We found a crossover tissue size above which both models coincide. Importantly, below this crossover size, the finite-domain model becomes a better approximation.

## Results

### 2.1. The reaction-diffusion model in the infinite domain

Here we briefly summarize the well-known reaction-diffusion model assuming an infinite domain and its analytic solution [10,11]. Within this model, it is assumed that the dynamics of the morphogen are faster than the proliferation rate of the tissue cells and, as a consequence, advective effects can be neglected. Otherwise, an advective term could be included to the model [17]. Since during developmental process tissues usually organize along a particular axis [18,19], this model is studied in a one dimensional setting [10,11]. It is assumed that the morphogen concentration *C*_1_(*x, t*) can diffuse with a diffusion coefficient *D* and is linearly degraded with a rate *k*.

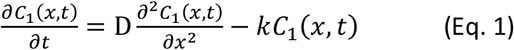

It is considered that there is no morphogen at the beginning, that is, the initial condition is:

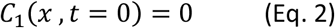

The only source of morphogen is a constant flux *q* located at the origin, represented by the first boundary condition:

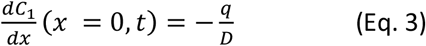

In this model, it can be assumed that there is a sink in the tip of the tissue absorbing the morphogen and assumes that the spatial domain is infinite:

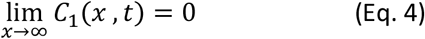

This model given by Eqs. 1-4 was extensively investigated by other authors, and the solution is [10,11]:

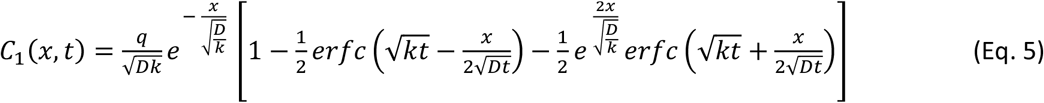

Where *erfc* (*x*) is the complementary error function.

Space and time variables can be rewritten in terms of the following dimensionless variables 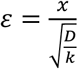 and *τ* = *kt*. Consequently, the morphogen flux at the tissue origin can be rewritten as 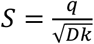 and the concentration as 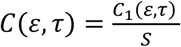 With this nondimensionalization, model equations (Eq.1-4) take the form:

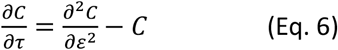

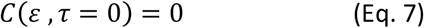

Where the morphogen source at the tissue origin, in nondimensional units, *ε* = 0, is:

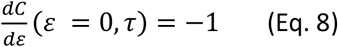

And a morphogen sink at infinite in the nondimensionalized units is now:

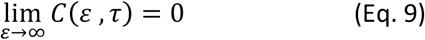

Which leads to this solution:

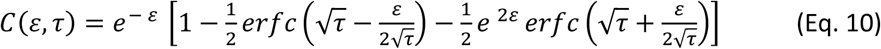

### 2.2. The reaction-diffusion model in finite domains: an analytical solution

The previous model variant entails an infinite domain (Eqs. 4 and 9). Since biological tissue sizes require a finite domain, we decided to replace the condition imposed by Eq. 4 with:

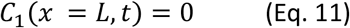

Where *L* is the length of the tissue. To our knowledge, the general solution for any given *L* is yet unknown.

We defined the quantity 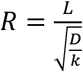, which is the only model parameter. This quantity represents the tissue length *L* in units of the characteristic reaction-diffusion length *λ*, defined as 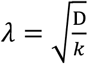 [20,21]. Thus, the second boundary condition for this model in nondimensionalized units is:

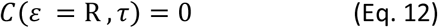

This equation replaces Eq. 9 in the section 2.1 assuming the finitude of the tissue.

We found the analytical solution of the general model for finite tissues (Eqs. 6-8 and 12) in the nondimensionalized units to be as follows (see Supplementary information for the demonstration):

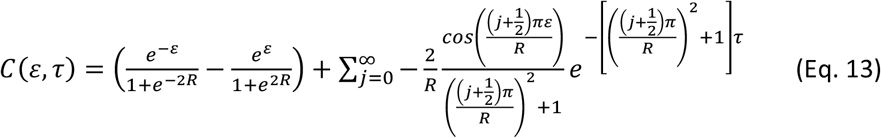

Moreover, we also found the solution for different boundary conditions such as assuming a non-null flux in *ε* = 0 and a zero flux in *ε* = *R* as well as a fixed non-null concentration in *ε* = 0 and a null concentration in *ε* = *R* (see Supplementary information). To further corroborate the analytical solution, we implemented the model numerically, by using a finite differences scheme (see Supplementary information). Our results indicate that the analytical solution is in agreement with numerical simulations (Fig. 1 in Supplementary information).

**Figure 1.**
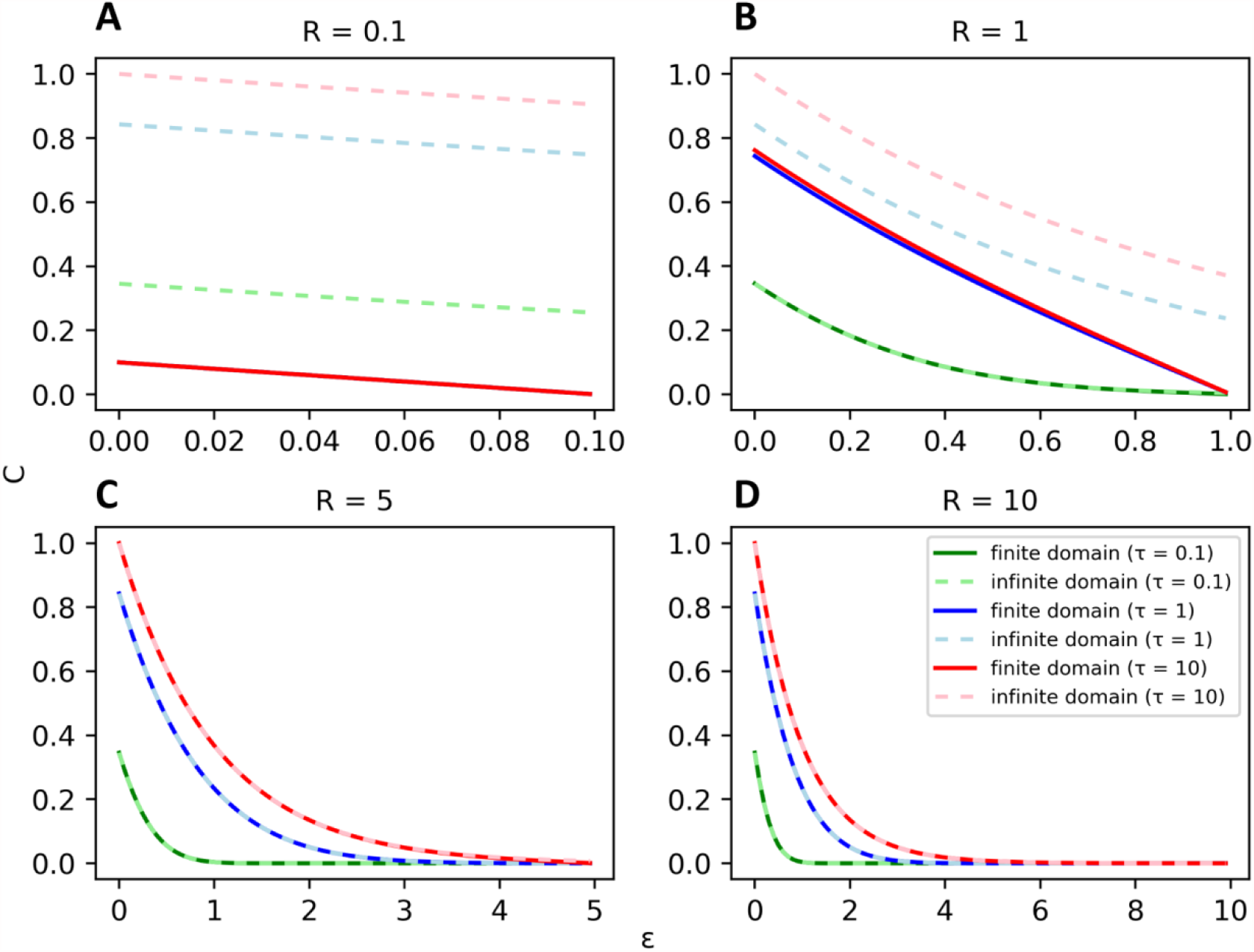
The morphogen spatial profile predicted from the reaction-diffusion model assuming finite domains at lengths larger than the crossover size converges to the profile predicted from the model assuming an infinite domain. Morphogen spatial profiles of the reaction-diffusion model assuming finite domains considering that the normalized tissue sizes *R* are **A)** 0.1, **B)** 1, **C)** 5 and **D)** 10, respectively, are depicted at three different times *τ* = 0.1, 1 and 10 (solid lines). The profiles from the model assuming infinite domains are also shown at the same times (dashed lines). *C, ε* and *τ* represent the normalized morphogen concentration, space and time, respectively. In panel **A)**, all spatial concentration profiles for finite domain overlap.

### 2.3 Transient morphogen distributions are qualitatively different between the model of infinite domain and the model of finite domains when they are of the order of the characteristic length *λ* or smaller

We decided to compare the reaction-diffusion model assuming a finite tissue *versus* an infinite domain. With the selected nondimensionalization, the latter does not have any free parameters. In contrast, the finite model has only one free parameter, *R*, which represents the tissue size in units of the characteristic length of the morphogen profile *λ*. By using our analytical solution for the model of finite tissues (Eq. 13), we explored the predicted morphogen spatial profiles at different tissue sizes (*i*.*e*., varying *R*) and compared them with those calculated from the previously known solution assuming an infinite domain (Eq. 10), at three different time points (Fig. 1). We observed that the morphogen concentrations predicted by the model assuming an infinite domain are higher than those predicted by the model assuming a finite domain (Fig. 1 A, B). For large enough tissue lengths, morphogen profiles predicted by both models are indistinguishable at each time point, as expected (Fig. 1 C and Fig 1 D). Hence, the previously reported model assuming an infinite domain is a reasonable description of the dynamics of morphogen profiles for larger tissues. However, when addressing a tissue whose length is of the order of the characteristic length *λ* or smaller, the model introduced in the present work is a more accurate description.

Moreover, we observed that large tissues lead to morphogen spatial distributions temporarily separated. In contrast, spatial distributions at different time points are indistinguishable in shorter tissues, suggesting that they already approached the steady state (Fig. 1A). This result would indicate that the larger the tissue, the longer the time necessary to reach the morphogen spatial distribution at the steady state (see also sections 2.4 and 2.6).

### 2.4. Steady state morphogen spatial distributions

The morphogen spatial distribution assuming an infinite domain at the steady state 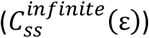 is well known [10,11] and with our nondimensionalization it is the following exponential spatial decay:

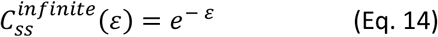

We calculated the steady state solution for our model of finite tissues, 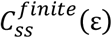, (Supplementary information):

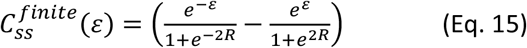

Increasing the tissue size in this model modifies the steady state profile, augmenting the maximum concentration at the origin and leading to a transition from a linear to an exponential curve (Fig. 2), in agreement with the results observed at any time (Fig. 1). Precisely, to estimate the limit when the tissue size tends to zero, we calculated the Taylor series expansion of the steady state solution (Eq. 15) on *R* to the first order. As *ε* is constrained by *R*, we subsequently obtained the Taylor series expansion for the resulting expression on *ε* to the first order:

**Figure 2.**
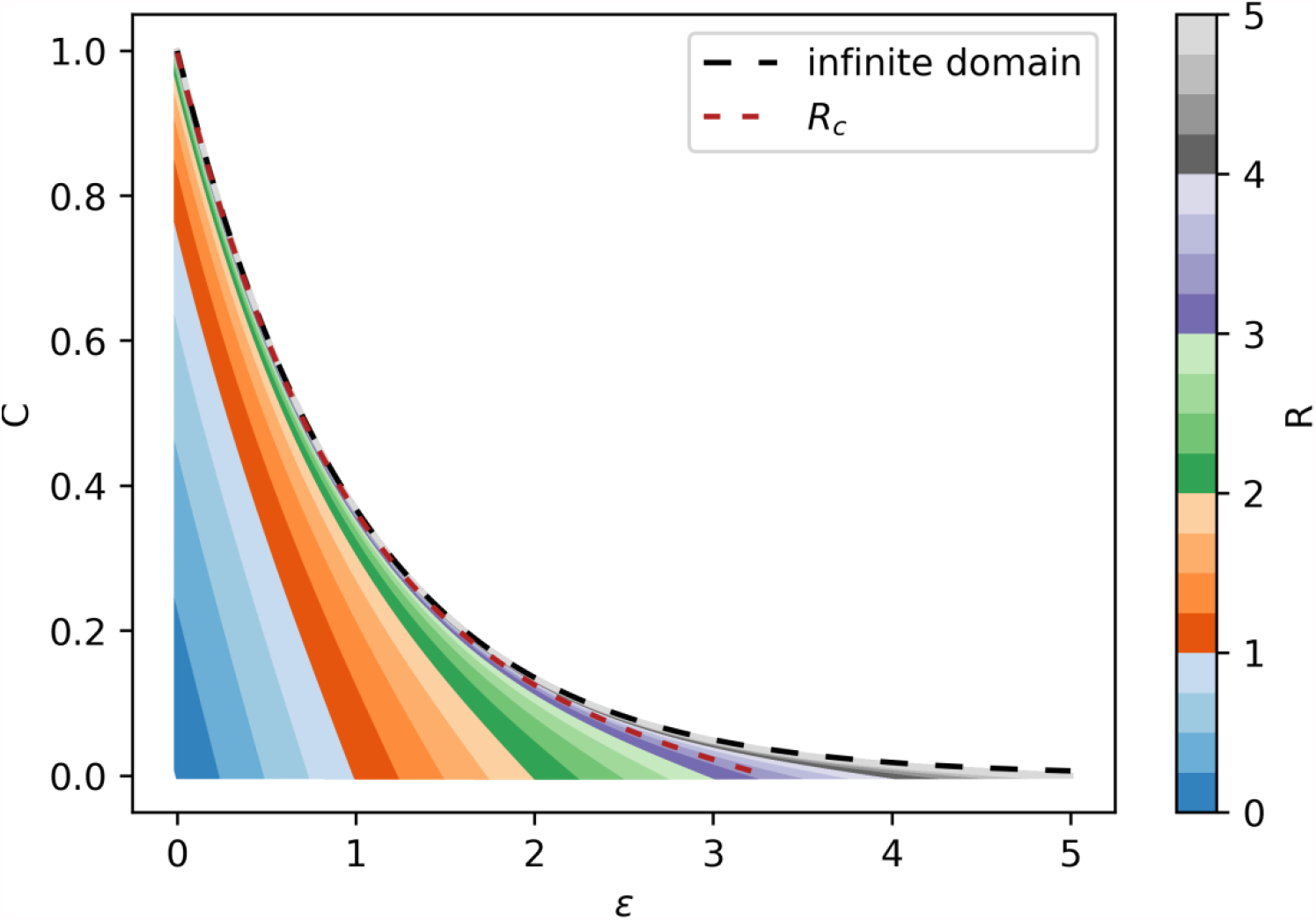
Morphogen spatial profile of the reaction-diffusion model assuming finite domains at the steady state transitions from a line-like to exponential-like spatial profile when the tissue size increases. Steady state profiles predicted from the reaction-diffusion model at different tissue sizes (*R*) are depicted. The steady state profile from the model assuming infinite domains and the crossover tissue size *R*_*c*_ (defined in section 2.5) are shown as dashed black and red lines, respectively.

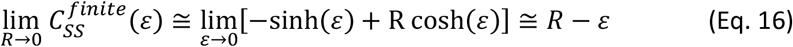

The limit when the tissue size tends to infinite was calculated:

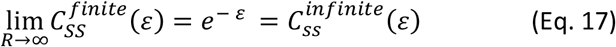

Remarkably, the steady state morphogen distribution of the finite model converges to the exponential distribution predicted by the infinite domain when the tissue length tends to infinity. Furthermore, by comparing the steady state solution (Eq. 15) with its complete solution (Eq. 13), we can re-write Eq. 13 as follows:

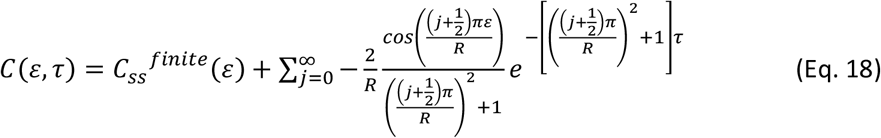

Where the second term of Eq. 18 vanishes when the time *τ* tends to infinity. Therefore, the morphogen concentration can be expressed as the steady state solution plus a term that describes a transient contribution.

### 2.5 Geometrical characterization of the morphogen spatial distributions

The steady state profiles predicted by the model of finite tissues changed from linear to exponential when increasing the tissue size as shown in section 2.4. In order to geometrically characterize the shape of the spatial profiles in the steady state regime, we defined *ε*_10_ as the dimensionless spatial position *ε* in which the morphogen concentration is 10 % of the concentration at the origin. When using this definition in the model assuming infinite domains, we obtain (see Supplementary information for details):

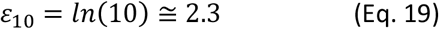

While for the model of finite tissues (see Supplementary information for details):

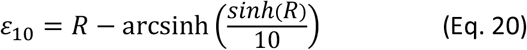

Thus, in the limit of small tissues, *ε*_10_ shows a linear dependence with the tissue size. However, when the tissue tends to infinity, *ε*_10_ becomes independent of the precise tissue size, reaching a plateau (Fig. 3). Additionally, when tissue size tends to infinity, in Eq. 20, *ε*_10_ recovers the value from the infinite model calculated in Eq. 19.

**Figure 3.**
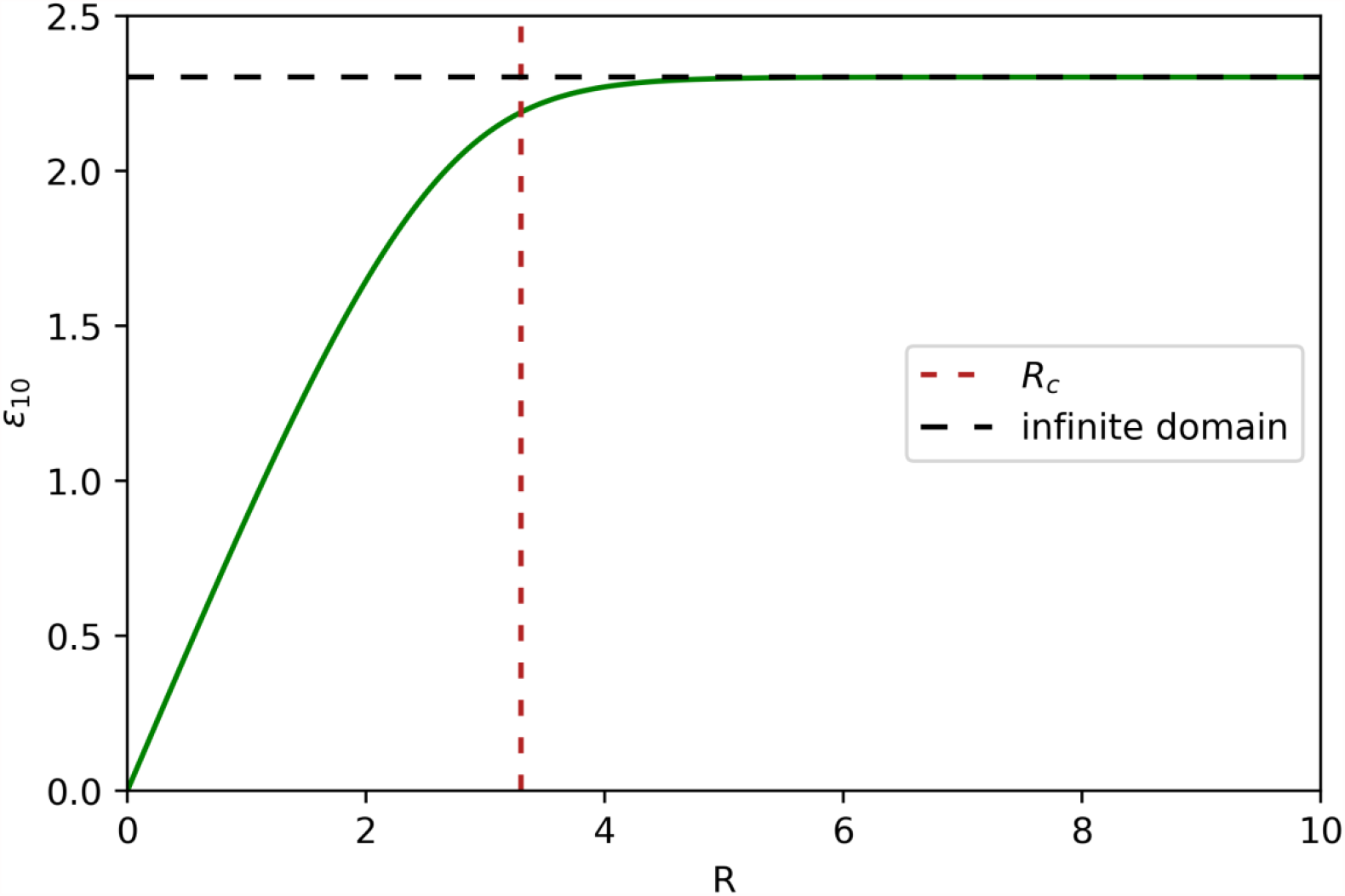
The geometrical factor *ε*_10_ characterizing the steady state spatial profiles of the reaction-diffusion model assuming finite domains (solid line) linearly grows with the tissue size (*R*, tissue length in nondimensional units) until it converges to the *ε*_10_ predicted from the model assuming an infinite domain (horizontal dashed black line). *ε*_10_ is defined as the spatial position (*ε*) where the morphogen concentration is 10 % of its value at the origin. The vertical dashed red line shows the crossover tissue size *R*_*c*_.

We wonder whether it is possible to establish a cut-off size to distinguish both regimes. To answer this question, we explore under what conditions, the shape of the morphogen spatial distribution depends on the tissue size. More precisely, we asked under what crossover tissue size *R*_*c*_ the geometrical observable *ε*_*10*_ would transition from linearly depending on the tissue size to becoming independent of it. To this end, we Taylor-expanded *ε*_*10*_ and arbitrarily looked for the *R* = *R*_*c*_ upon which the second non-zero term of the series would be about 20 % of the first linear term (See Supplementary information for details). Our results show that the crossover tissue size separating both regimes is about 3 times the characteristic length *λ* (*R*_*c*_ ≈ 3.3).

The analysis of the dependency of *ε*_*10*_ with the tissue size can also be made before the morphogen distribution achieves the steady state. Although we could not find an analytical expression for this observable in the general case, we explored this dependency numerically (Fig. 4). We observed that, for each tissue size, *ε*_*10*_ increases in time until it reaches a plateau, which indicates that the spatial profile stabilizes in the steady state. Moreover, the time needed to reach the plateau monotonically increases with the tissue size until *R* ∼ *R*_*c*_. For larger tissue sizes, the time to reach the plateau converges to the prediction of the model for infinite domains (Fig. 4). This result is consistent with the fact that morphogen spatial distributions at different times are overlapped in smaller tissues and separated in larger ones (Fig. 1). This is a consequence of the second term of Eq. 18: the larger the tissue, the longer waiting times are required to vanish the exponential in the second term.

**Figure 4.**
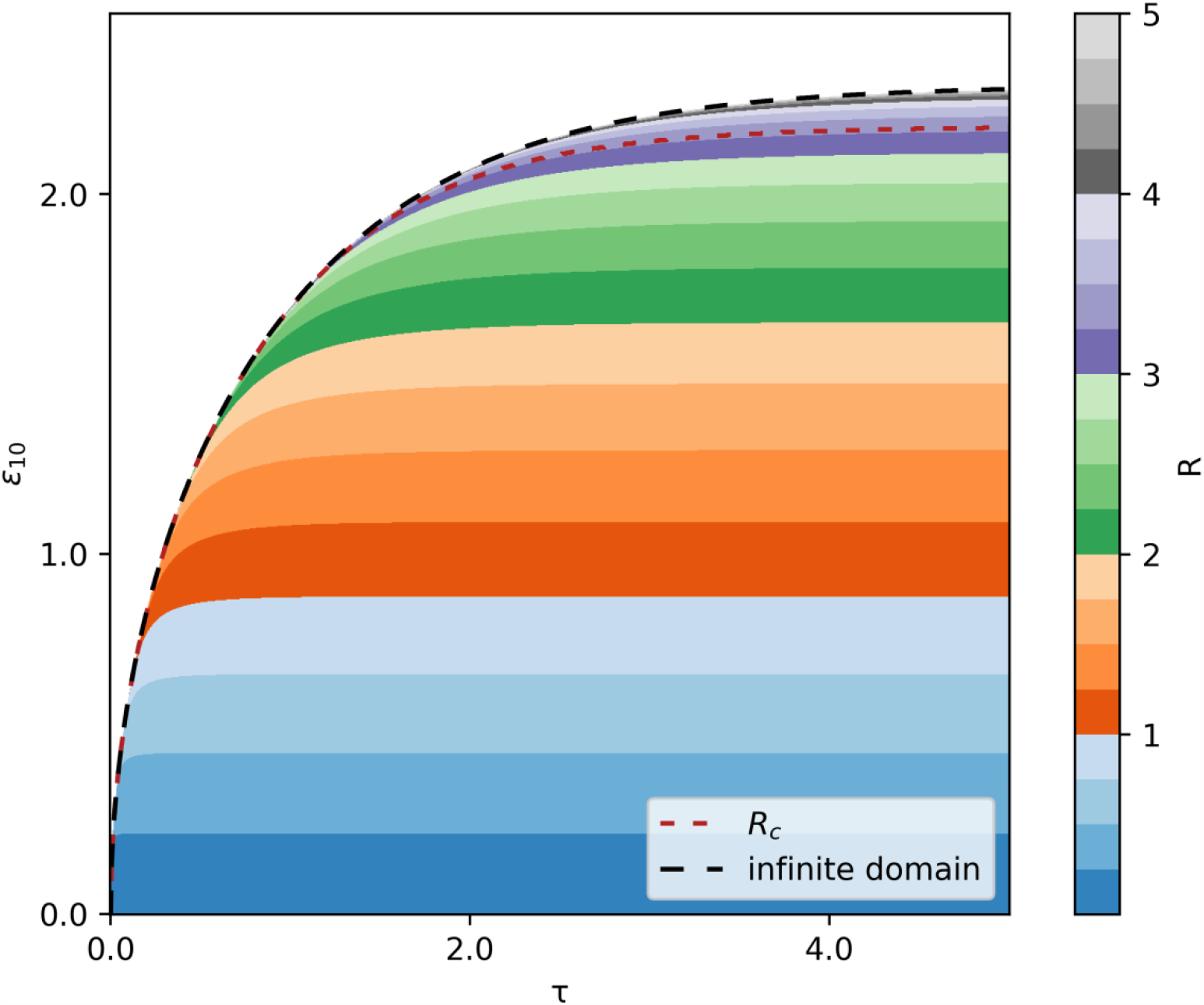
The larger the tissue, the longer the time to reach the steady state. Kinetics of the geometrical factor *ε*_10_ predicted from the model for finite tissues of different sizes (*R*, tissue length in normalized units). The kinetic of the factor *ε*_10_ of the model for an infinite domain is also shown (dashed black line). *ε*_10_ is defined as the spatial position (*ε*) where the morphogen concentration is 10 % of its value at the origin. The dashed red line shows the crossover value *R*_*c*_.

### 2.6 Time to reach the steady state morphogen distribution

In Section 2.5 we suggested that the larger the tissue, the longer it takes the model to reach the steady state. To test this hypothesis, we took advantage of a method developed by Berezhkovskii and colleagues [11] to quantify the mean time (*μ*_*τ*_) it takes a morphogen profile to reach its steady state. They applied this method to the reaction-diffusion model assuming an infinite domain and obtained (in our nondimensional units):

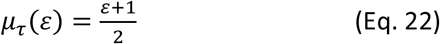

That is, the mean time to reach the steady state is linear with the position within the infinite domain. We applied the same method to our reaction-diffusion model of finite tissues and obtained (see Supplementary information for details):

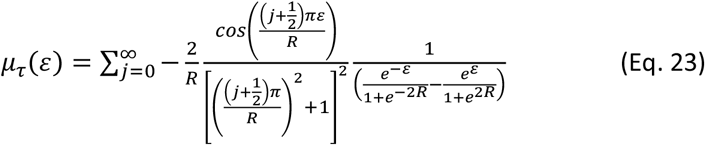

Thus, for our model, the mean time to reach the steady state not only depends on the position within the tissue but also on the tissue size.

To formally compare the mean times calculated from both reaction-diffusion models we also need to estimate a measure of the error. Hence, we calculated the standard deviation of the time to reach the steady state, *σ*_*τ*_ (see Supplementary information for details). For the model assuming an infinite domain, it reads:

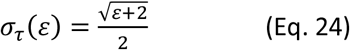

Which coincide with the already reported result by Ellery and colleagues [15]. In contrast, for the reaction-diffusion model for finite tissues, we obtain:

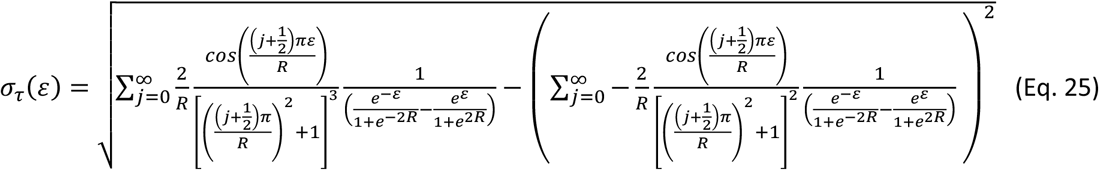

As with the mean, the standard deviation of the time necessary to reach the steady state not only depends on the positions along the tissue but also on the tissue size. At the origin of the tissue (*ε* = 0), both, *μ*_*τ*_ and *σ*_*τ*_, increase with *R* until they converge toward 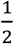 and 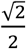, respectively, when *R* tends to infinite (Fig. 5 A). These are precisely the expected values from the model assuming infinite domains evaluated at the tissue origin (Eqs. 22 and 24). Interestingly, the transition between the domains in which *μ*_*τ*_ and *σ*_*τ*_ depend on the tissue size and where they are independent of it coincides with the crossover tissue size of about 3 *λ* determined in the previous section (compare Fig. 5 A with Fig. 3). The ratio between them, constituting the Coefficient of Variation 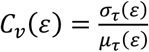 also experiences a transition near the crossover tissue size until converging to 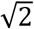 (Fig. 5 A).

**Figure 5.**
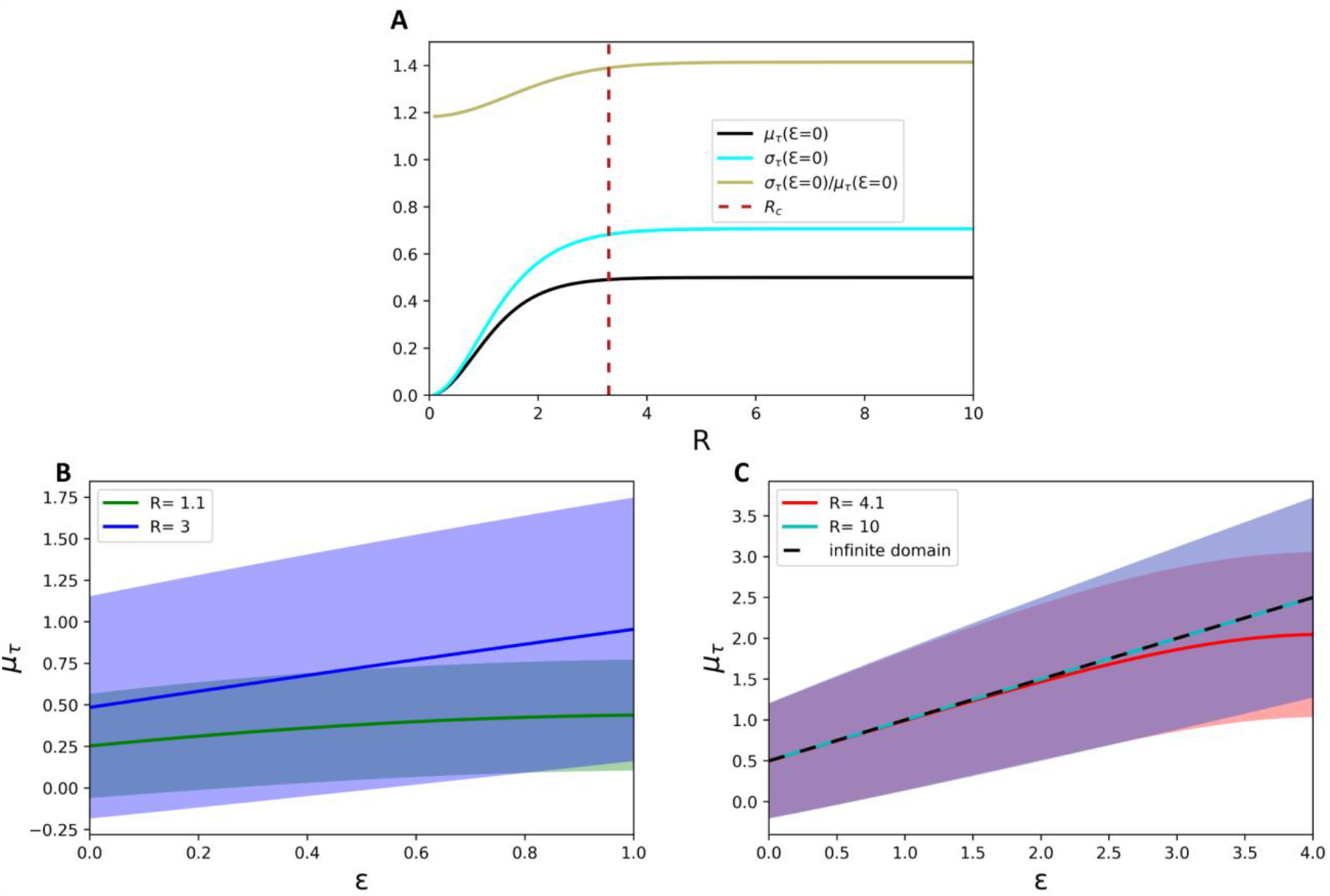
Crossover tissue size separates two regimes of the time to reach the steady state. **A)** The mean time to reach the steady state (black line), the standard deviation (light blue line) and the variation coefficient (brown line) as a function of *R* predicted from the model assuming finite domains at *ε* = 0. The vertical dashed red line indicates the crossover tissue size *R*_*c*_. **B)** Spatial profile of the mean time to reach the steady state for *R* = 1.1 (green line) and *R* = 3 (blue line). The standard deviation (*σ*_*τ*_) is represented by the shady areas surrounding the curves. **C)** Spatial profile of the mean time to reach the steady state for *R* = 5.1 (red line) and *R* = 10 (light blue line). The result for the model of an infinite domain is shown in dashed black line. The standard deviation (*σ*_*τ*_) is represented by the shady areas surrounding the curves.

For tissues smaller than the crossover size, the mean time to achieve the steady state and its error in each position strongly depend on tissue size (Fig. 5 B). On the contrary, for tissue sizes higher than the crossover tissue size, both magnitudes become independent of the size (Fig. 5 C). Importantly, for tissues smaller than the crossover size, the steady state will be reached significantly faster than the prediction from the model assuming an infinite domain. For tissues larger than the crossover size, both models agree in the time to achieve steady state (Fig. 5 B and C).

### 2.7 Finite *versus* infinite domains in the reaction-diffusion model used in the FRAP-based determination of diffusion parameters

Diffusion parameters of morphogens can be experimentally determined in tissues by using Fluorescence recovery after photobleaching (FRAP) experiments [22, 23]. From this technique, the diffusion coefficient *D* and degradation constant *k* are obtained indirectly by fitting to experimental concentration measurements the analytical solution of the model assuming a finite domain [23,34] as well as an infinite domain [19, 24]. Thus, we wondered whether the election of the model used in FRAP has an impact on the calculated *D* and *k* values. To that end, as a proof of principle, we evaluated whether the infinite domain model could render an accurate estimation of the kinetic parameters D and k, when fitted to a dataset simulated with the finite domain model used as a proxy for experimental data. We simulated steady state concentration profiles by using the finite-domain model (Eq. 15) rewriting the concentration in the original coordinate *x* = *λε* and arbitrarily setting *λ* = 1 for different values of *L*. Then, we rewrote the steady state concentration of the infinite-domain model (Eq. 14) in the original coordinate *x* and performed a curve fitting for each of the datasets obtained using Eq. 15. We used Eq. 14 in the original coordinate as fitting function and *λ* as the free parameter. We obtained the predicted value of *λ* as a function of *R* = *L* (Fig. 6). For large values of *R*, the predicted *λ* is approximately 1, which is in agreement with the value actually used to generate the data. In contrast, for values of *R* smaller than *R*_*c*_, the predicted value of *λ* deviate from 1, converging to 0 for small values of *R*. We concluded that both models can be used to infer the kinetic parameters *D* and *k* from FRAP experiments, provided that tissue sizes are higher than *R*_*c*_. On the contrary, for tissues smaller than this crossover value, the model assuming finite domains is the best alternative.

**Figure 6.**
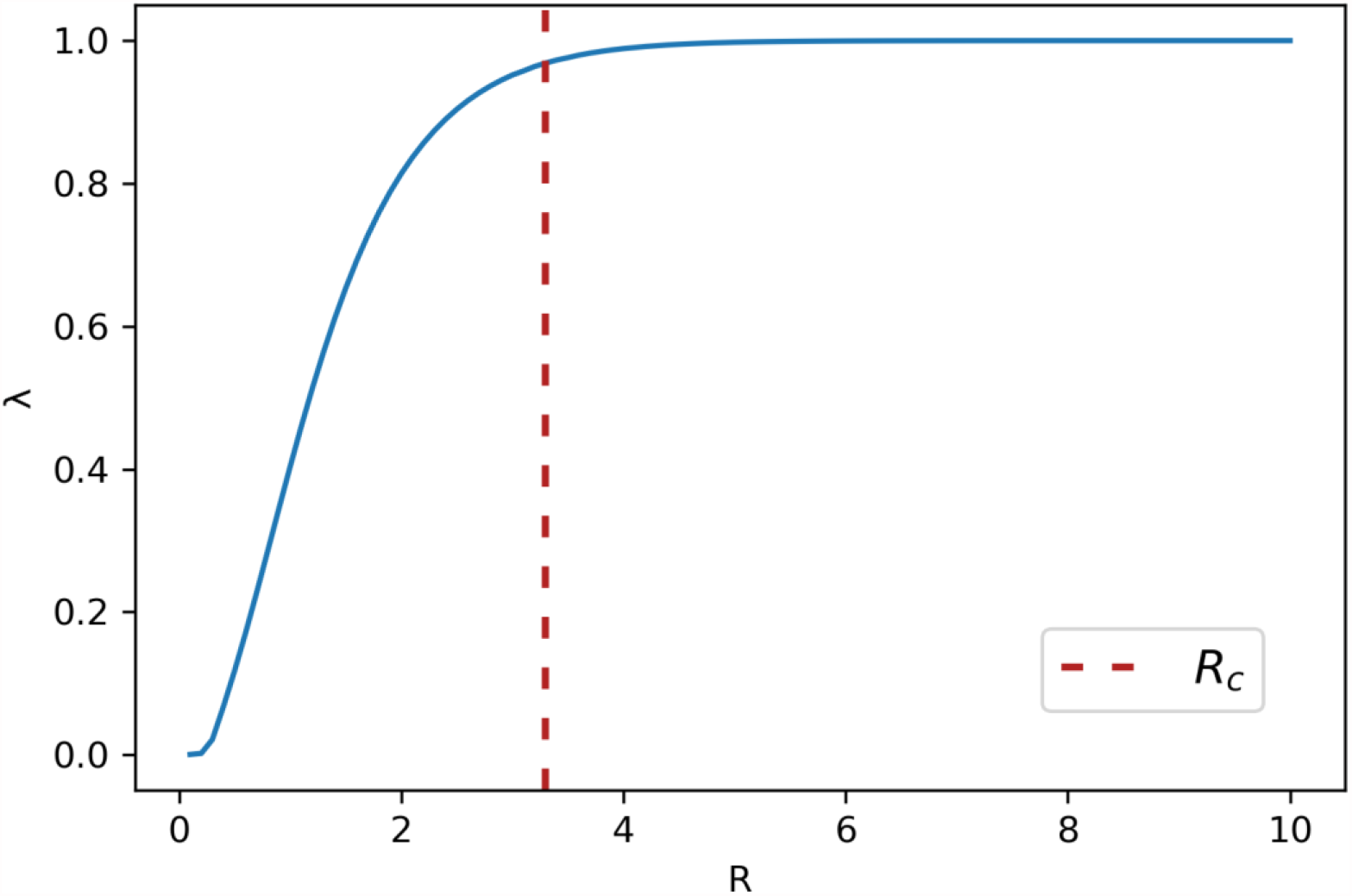
The reaction-diffusion model assuming an infinite domain does not correctly predict the characteristic length for tissues smaller than the crossover size. The concentration of a morphogen in steady state was simulated with the model assuming finite domains along 101 equidistant positions from 0 to *L*, for different values of *L* going from 0.1 to 10. *λ* was arbitrarily set equal to 1. The simulated curves were fitted with the model assuming an infinite domain where *λ* was the only free fitting parameter. For large values of *R* the infinite domain model predicts correctly the value *λ* = 1. For *R* smaller than *R*_*c*_ (depicted by the vertical dashed red line) the predicted *λ* goes to 0.

## Discussion

Reaction-diffusion models were conceived in the seminal article of Alan Turing to hypothesize under what conditions heterogeneous patterns could emerge from a homogeneous one in tissue morphogenesis [1]. After the concept of positional information was posed by Lewis Wolpert [25], as illustrated by his well-known French Flag Problem ([26]; see also the review by Sharpe [27]), reaction-diffusion models resurfaced to account for mechanisms capable of generating spatial gradients that could serve as positional signals. Francis Crick was entertaining the hypothesis of reaction-diffusion signals as probable morphogenetic driving forces [28]. Reaction-diffusion models were specifically studied by Alfred Gierer and Hans Meinhardt to understand pattern formation in tissue development and regeneration [2]. Thereafter, a plethora of reaction-diffusion models were developed and proposed over the years to describe different morphogen gradients [29,30,31,32]. Some notable examples are Bcd in the syncytial *Drosophila* embryo [33], Dpp in developing wing imaginal disc in *Drosophila* [19], Fgf8 in the gastrulating *Danio rerio* embryo [34], among other examples. Despite the controversy of whether reaction-diffusion models represent an effective or accurate description of tissue pattern formation, these modelling framework became an essential construct to guide mathematical approaches in development [5,35] and regeneration [36].

In this study, we investigated the spatiotemporal distribution of a morphogen with a minimal reaction-diffusion model in a finite domain, as a proxy for a tissue. The solution of the model assuming an infinite domain has been already reported [10,11]. A number of reaction-diffusion models were previously considered to investigate morphogen gradients in finite domains, by means of numerical simulations (see, for instance, [7,12], among other examples). A reaction-diffusion model assuming finite domains was exactly solved assuming Neumann boundary conditions to investigate scaling of morphogens in tissues [13] and robustness of pattern formation in development [14], among other examples. A similar model was considered to investigate cell migration and proliferation of a population of precursor cells on a uniformly growing tissue by Simpson [37], based on the model of cell colonization in uniformly growing domains [38]. In his model, Simpson [37] explored a more general case of a growing domain, which can recapitulate the case of a fixed domain by setting the growth speed to zero. Nevertheless, because the model focused on cells instead of morphogens, it assumed a positive reaction term to account for cell proliferation and a non-zero initial condition, in contrast to our negative reaction term and our zero initial condition. Hence, imposing a zero initial condition in this previously reported model yields the null solution.

The analytical solution here reported could be instrumental in computational packages devoted to multi-scale modelling, which involve a signalling scale coupled with a cellular scale. Although their cellular layer could entail a Cellular Potts Model (CPM) [39, 40] in CompuCell3D [41] and MORPHEUS [42], or a vertex model [43, 44] in CHASTE [45], their signalling scale is typically modelled by a reaction-diffusion scheme. Since in these packages a finite domain is the only possible choice, they cannot avoid a numerical implementation. While our numerical results, based on a finite-difference algorithm cannot be distinguished from the analytical solution (Fig. 1 in Supplementary information), the last one is naturally more accurate and computationally more efficient (see Supplementary information), which could prove useful for multi-scale modelling implementations. Likewise, this new solution could help to improve the calculation of recovery curves in FRAP experiments, as for tissues below the crossover size *R*_*c*_, the model assuming finite domains is a better approximation.

Our results showed that the morphogen spatial distributions predicted by our model assuming finite domains depend on the only relevant model parameter: the normalized tissue size *R*. By determining the spatial position along the tissue where the morphogen concentration is 10 % of the source concentration (*ε*_*10*_), we geometrically characterized the steady state spatial distribution. This characterization led us to find two regimes within the parameters space, separated by a crossover tissue size *R*_*c*_ (Fig. 3 and Fig. 7C and 7D). For tissues longer than *R*_*c*_, the distributions are exponential-like and cannot be distinguished from those predicted from the model assuming an infinite domain (Fig. 2 and Fig. 7B). In this regime of the parameter space, the mean and standard deviation of the time to reach the steady state (evaluated at the tissue origin) do not change much with the tissue size and converged towards the corresponding values from the model assuming an infinite domain (Fig. 5 and Fig. 7F). When comparing the morphogen concentrations predicted by both models we found that the difference between them is mostly negligible (Fig. 8 A and B). Hence, the model assuming an infinite domain can be considered a good approximation of the model assuming finite domains for tissue sizes larger than *R*_*c*_.

**Figure 7.**
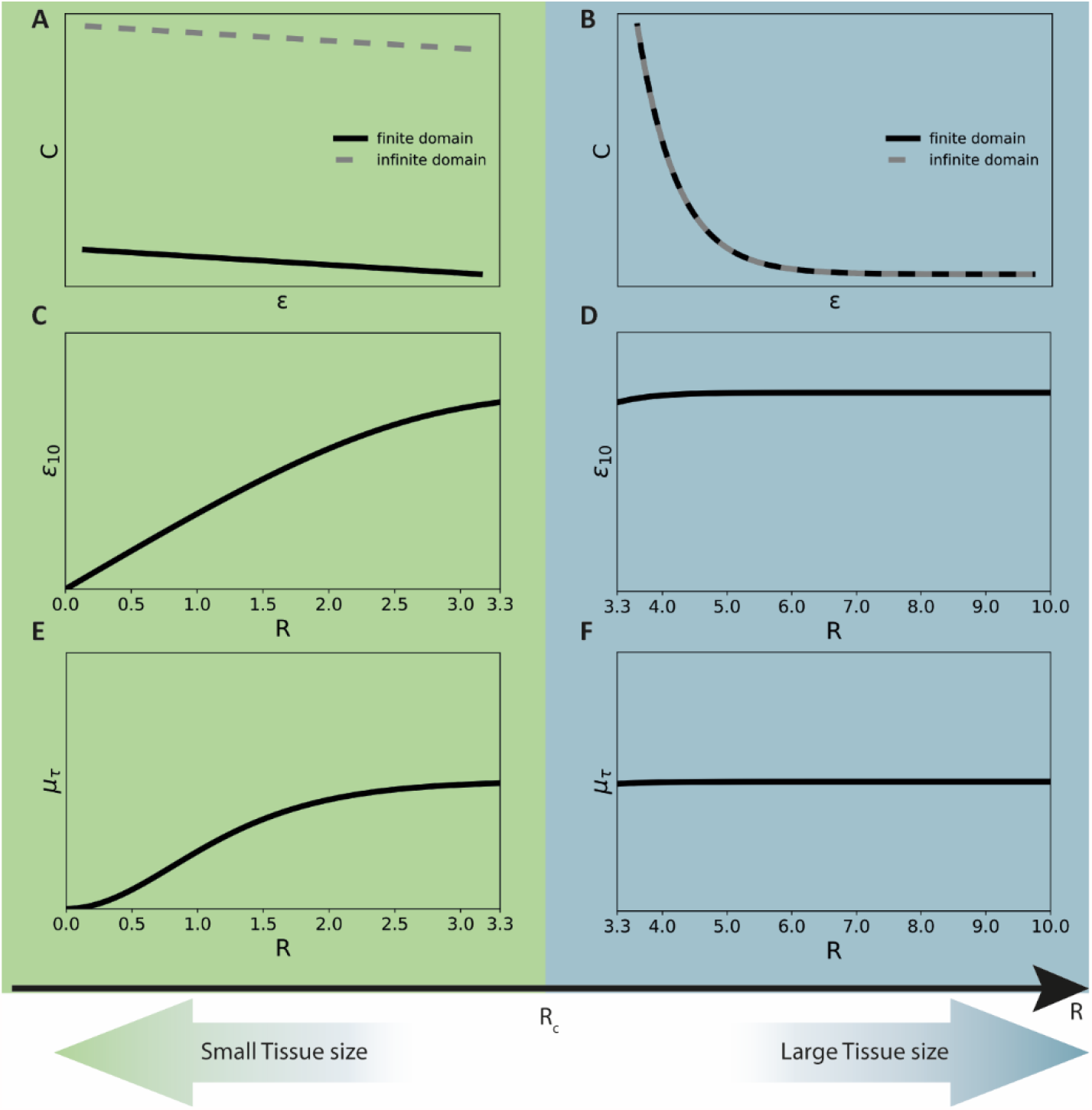
Transition between small and large tissues: Two reaction-diffusion regimes separated by a crossover tissue size. Sketch summarizing the main differences between small and large tissue sizes separated by a crossover tissue size (*R*_*c*_). Spatial profiles of morphogen concentration *C* **(A, B)** and dependency of the geometrical factor *ε*_*10*_ **(C, D)** and the mean time to reach the steady state *μ*_*τ*_ **(E, F)** with the nondimensionalized tissue size *R*, for small (A, C, E) and large tissues (B, D, F).

**Figure 8.**
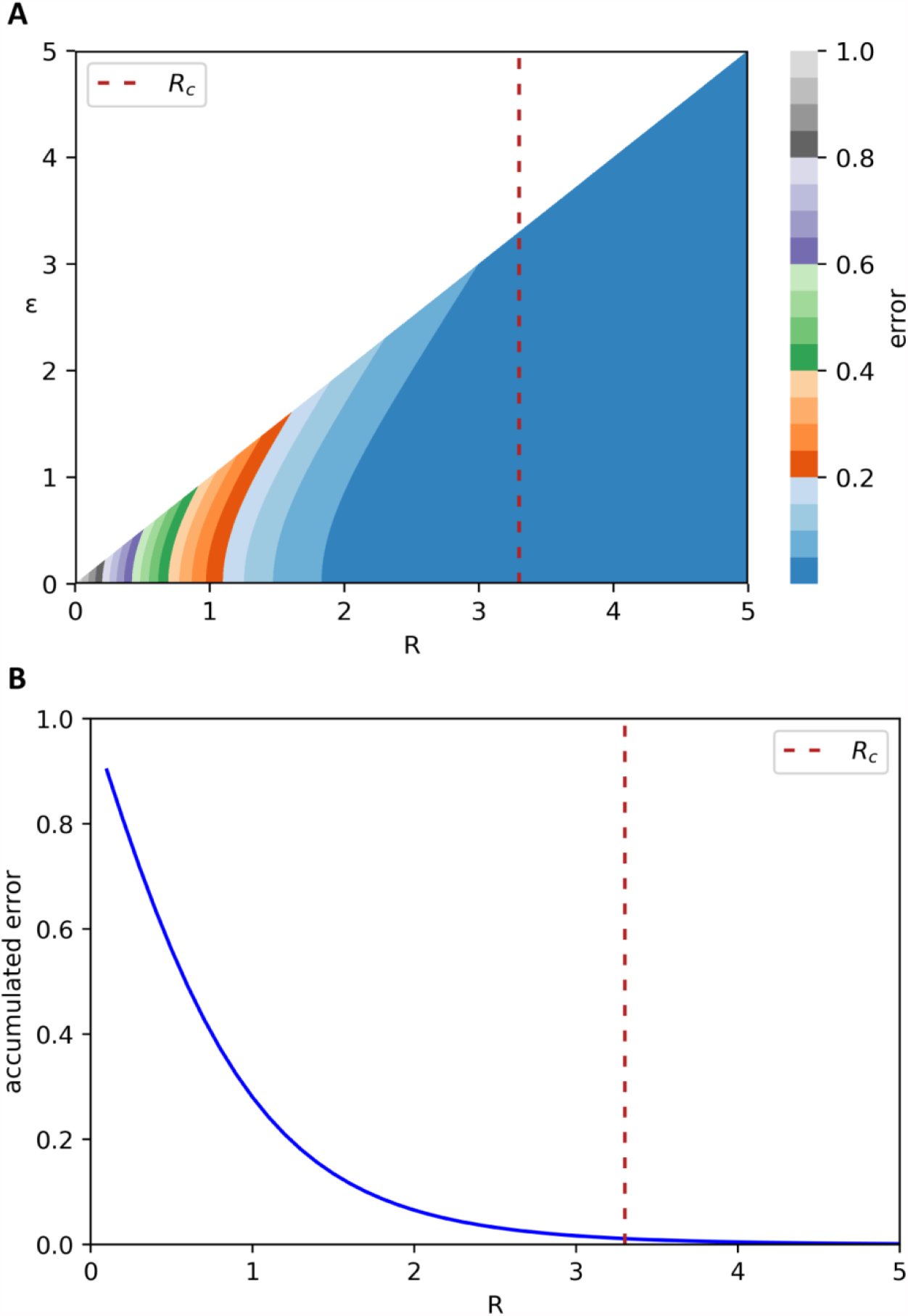
The reaction-diffusion model assuming a finite domain is a better approximation than the model assuming an infinite domain when the tissue size is smaller than the crossover tissue size. **A)** Heat map showing the difference of morphogen concentration predicted from the reaction-diffusion model assuming a finite and the infinite domain as a function of the position within the tissue (*ε*) and the tissue size (*R*). This difference could be considered as the error committed when utilizing the standard model assuming the infinite domain at a given position *ε* from a tissue of size *R*. **B)** The difference calculated in (A) integrated over the tissue and normalized with *R*, as a function of *R*, representing the global error of using the standard model assuming the infinite domain when the tissue size is *R*. The vertical dashed line indicates the crossover tissue size *R*_*c*_.

In contrast, for tissues smaller than *R*_*c*_, the distributions tend to be linear and are clearly separated from those predicted with the model assuming an infinite domain (Fig. 2 and Fig. 7A). Furthermore, the time to reach the steady state strongly depends on the tissue size in this regime (Fig. 5 and Fig. 7E). In particular, the error of using the model assuming an infinite domain increases when *ε* tends to *R* and the smaller the tissue the higher the error accumulated over the entire tissue (Fig. 8 A and B) (See Supplementary information for details). Thus, our results indicate that to investigate tissues smaller than approximately 3 times the characteristic length *λ*, the model assuming finite domains should be used.

The crossover tissue size provides a straightforward criterion to decide when to use any of the two models presented here. As an example, the characteristic length of Wg was estimated in 6 μm in the *Drosophila* wing disc, where the tissue size was about 50 μm [19]. The resulting *R* ∼ 8 > *R*_*c*_ indicates that the model assuming an infinite domain is a reasonable approximation in this scenario. A similar conclusion can be drawn when studying Dpp in the *Drosophila* haltere. For this morphogen, the characteristic length and the tissue size can be estimated in ∼ 10 and ∼ 100 μm, respectively [7], which leads to *R* ∼ 10 > *R*_*c*_. In contrast, the last morphogen, Dpp, but in the *Drosophila* wing disc has a characteristic length of 20 μm [19] which implies *R* ∼ 2.5 < *R*_*c*_. As a consequence, the model assuming finite tissues is the most correct approximation to describe morphogen propagation in this scenario. Something similar occurs with Fgf8 in the *Danio rerio* embryo, whose characteristic length was estimated as 200 μm while the tissue size is about 200 μm [34], from which a *R* ∼ 1 < *R*_*c*_ can be calculated. By only looking at the previous examples, it is clear that there is no correlation between the model selection and the morphogen under study, since the same morphogen, Dpp dynamics is better explained with the model assuming finite domains in the *Drosophila* wing disc while in the *Drosophila* haltere the model assuming an infinite domain is actually sufficient. The same lack of correlation can be observed between the model selection and the tissue of interest. Indeed, in the same tissue, *Drosophila* imaginal disc, Wg could be described with the model assuming an infinite domain while Dpp requires the most precise model of finite domains.

In conclusion, we found two reaction-diffusion regimes for large and small tissues, separated by a crossover tissue size. While above this crossover the infinite-domain model constitutes a good approximation, it breaks below this crossover, whereas the finite-domain model faithfully describes the entire parameter space. Further studies will be needed to unveil the spatiotemporal distribution of morphogens in tissues whose size is not fixed. Our finding of the delineated crossover tissue size could be instrumental to select the proper reaction-diffusion model in future studies aimed to address tissue morphogenesis and other relevant problems regarding pattern formation in biology and medicine.

## Computational methods

In this article, the reaction-diffusion model assuming a finite domain and its comparison with the model assuming an infinite domain were studied. The analytical derivation of the reaction-diffusion model assuming a finite domain for different boundary conditions is presented in the section 1 in Supplementary information. Comparison between analytical and numerical solutions is described in the section 2 in Supplementary information. Steady state calculations, the geometrical characterization of the spatial distribution profiles given by *ε*_10_ and the estimation of the crossover tissue size *R*_*c*_ are shown in the sections 3, 4 and 5 in Supplementary information, respectively. Mean time to reach the steady state together with its standard deviation are in the section 6 in Supplementary information. Details on the error of assuming an infinite domain instead of a finite domain in the steady state solutions are in the section 7 in Supplementary information. Finally, the efficiency of the analytical solution *versus* the numerical one is analysed in the section 8 in Supplementary information.

All model calculations were encoded in Python 3.7.3 and performed using NumPy [46] and SciPy [47] while visualization was executed with matplotlib [48] and seaborn [49]. The source codes for all the calculations and figures were implemented in supplementary notebooks using Jupyter Notebook (http://jupyter.org/) and can be found at: http://doi.org/10.5281/zenodo.4421327 [50].

## Acknowledgements

We thank Diane Peurichard and Valeria Caliaro from the INRIA Paris - team MAMBA and the Laboratoire Jacques Louis Lions (LJLL)-Sorbonne Université, Juan José Gervasio from the University of La Plata, Fabian Rost from the Center for Molecular and Cellular Bioengineering (CMCB), Technische Universität Dresden and the Sys*Bio* members of the Chara lab for their invaluable comments on this study.

